# LM-Merger: A workflow for merging logical models with an application to gene regulation

**DOI:** 10.1101/2024.09.13.612961

**Authors:** Luna Xingyu Li, Boris Aguilar, John H Gennari, Guangrong Qin

**Affiliations:** Institute for Systems Biology, Seattle, WA 98109, United States of America; Department of Biomedical Informatics and Medical Education, University of Washington, Seattle, WA 98195, United States of America

**Keywords:** gene regulatory networks, logical models, model integration, acute myeloid leukemia, systems biology

## Abstract

**Motivation:** Gene regulatory network (GRN) models provide mechanistic understanding of genetic interactions that regulate gene expression and, consequently, influence cellular behavior. Dysregulated gene expression plays a critical role in disease progression and treatment response, making GRN models a promising tool for precision medicine. While researchers have built many models to describe specific subsets of gene interactions, more comprehensive models that cover a broader range of genes are challenging to build. This necessitates the development of automated approaches for merging existing models.

**Results:** We present LM-Merger, a workflow for semi-automatically merging logical GRN models. The workflow consists of five main steps: (a) model identification, (b) model standardization and annotation, (c) model verification, (d) model merging, and (d) model evaluation. We demonstrate the feasibility and benefit of this workflow with two pairs of published models pertaining to acute myeloid leukemia (AML). The integrated models were able to retain the predictive accuracy of the original models, while expanding coverage of the biological system. Notably, when applied to a new dataset, the integrated models outperformed the individual models in predicting patient response. This study highlights the potential of logical model merging to advance systems biology research and our understanding of complex diseases.

**Availability and implementation:** The workflow and accompanying tools, including modules for model standardization, automated logical model merging, and evaluation, are available at https://github.com/IlyaLab/LogicModelMerger/.

## Introduction

Gene regulatory networks (GRN) are fundamental to a wide range of biological processes, including cell differentiation, signaling transduction, and cell cycle in both normal and disease states (1). GRNs provide a virtual representation of biological systems, which can then be used for simulation of the dynamics of the physical systems. With these, we can make predictions about disease progression or drug response in a personalized manner, taking into account an individual’s genetic profiles for genes included in the model. GRNs can be modeled using various approaches, including logical models, ordinary differential equations (ODEs), and piecewise linear differential equation models (2). Among them, logical models are appealing due to their simplicity and versatility, especially in cases where kinetic parameters are unavailable (3,4). Logical GRN models have been successfully applied to study a wide range of diseases, including breast cancer (5), pancreatic cancer (6), and acute myeloid leukemia (AML) (7).

Building GRN models requires extensive domain expertise. Due to the limitation of knowledge from individual research teams, and the focus of different studies, existing published models are usually focused on one process or theme, with limited coverage of genes and processes for a more systematic study. Different models also use different representations for the nodes and relationships, which further complicates any efforts at model integration. To address this, the community has emphasized the need for systematic methods to integrate previously developed GRNs (e.g., in (2,5)). Such integrated models are essential for illustrating complex biological phenomena and understanding the intricate regulatory mechanisms that drive cellular behavior (8,9).

One integration approach is to develop modular models, where distinct biological processes are modeled independently and combined during simulation via variable transformations and synchronization (10). While modular models hold promise, they face challenges in standardization and harmonization across models, which remain labor-intensive and error-prone. Furthermore, modular models often fail to preserve key network features, such as feedback loops and dynamic interactions among components, as their simulations are kept independent. Directly integrating individual models overcomes these limitations by preserving network structures and enabling continuous information exchange. Tools like RegNetwork (11) and NDEx (12) provide solutions by leveraging shared regulatory components from multiple sources, but they primarily focus on visualization and storage rather than computational analysis. Whole-cell models (WCM) represent another ambitious effort to simulate entire cellular processes by integrating diverse biological networks (13). Yet, WCMs are currently limited to a few organisms such as *M. genitalium* (14) and *E. coli* (15), and integrating heterogeneous data remains a major challenge (16). Notably, most existing tools and workflows focus on ODE and rule-based models, leaving logical models underserved. This underscores the need for a dedicated workflow to merge logical models effectively.

In this study, we developed a workflow for merging logical GRNs models (LM-Merger), and demonstrated its effectiveness in enhancing the understanding and coverage of GRNs. We applied this workflow to AML, a highly aggressive hematopoietic malignancy characterized by the accumulation of somatic genetic alterations that disrupt normal cell behavior (17–22). AML cells from different patients often carry diverse mutations (19,20,22,23), which are associated with varied drug responses (22). Numerous GRN models have been developed to investigate different aspects of AML, including hematopoietic stem cells regulation (24), potential drug responses (25), and predictions of clinical outcomes (7). Merging these models offers a promising approach to creating a more comprehensive GRN model for AML. Such an integrated model better captures the complexity of the disease, improves drug response predictions, and helps identify patient-specific therapeutic targets. By combining two pairs of published AML models, we created larger, more comprehensive models that retained the behaviors of original models while offering broader insights. These merged models are better suited for predicting gene expression and clinical outcomes in AML patients. Furthermore, this workflow is adaptable for applications in other logical models beyond.

## The Logic Model Merging Workflow

The LM-Merger workflow integrates logical models from various sources into a unified representation to provide a more comprehensive view of biological mechanisms. It includes five main steps: (1) Finding models, (2) Standardizing and annotating models, (3) Reproducing selected models, (4) Merging models, and (5) Evaluating the merged model (Fig 1A). While this section focuses on describing the workflow, details on its implementation in specific use cases are provided in the Supplementary Data.

**Fig 1:**
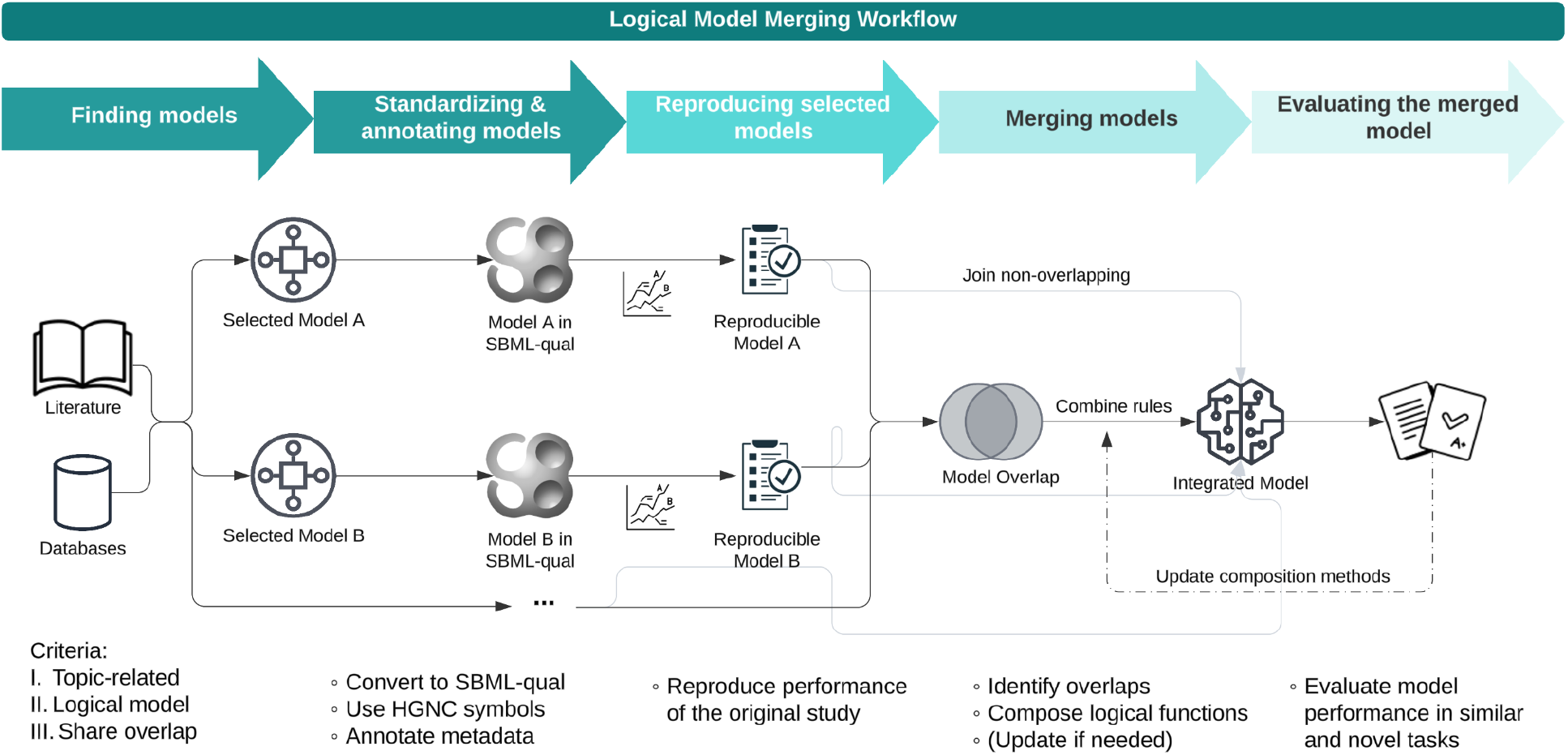
Overview of the logical model workflow. The flowchart illustrates the step-by-step process for merging logical models. The workflow begins with collecting eligible models from literature and databases. These models are then standardized into the SBML-qual format with added annotations. Reproducibility is verified for each model before proceeding to the merging step. The merging steps include composition of rules for overlapping nodes and non-overlapping components are directly integrated. Finally, the integrated mode is evaluated on tasks similar to their original studies and novel tasks of interest, ensuring its accuracy and applicability.

### Finding models

The initial phase of integrating logical models involves identifying candidate models. This typically starts with existing data like publications and repositories relevant to the biological system under investigation. Researchers often look into large repositories of logical models, such as the Cell Collective (26), the GINsim repository (27) and BioDiVinE (28), to find logical models of interest. Pathway databases like KEGG (29), Reactome (30), and causal interactions such as SIGNOR (31) can also be useful resources, although they only provide interaction graphs and need to be translated into logical relationships afterwards. A simple approach to this translation is to use the ‘Inhibitor Wins’ combination method, where inhibitory interactions dominate over activatory ones (See the following section).

After collecting logical models and building a library of candidate models, each model should be carefully reviewed to assess its relevance based on the topic, components involved, and the knowledge source. The decision should be primarily based on the purposes of the models and their biological relevance. Another important consideration is the level of overlap, as we aim for the models to complement each other by expanding the coverage of nodes while sharing key regulators. This ensures the integrated model is both comprehensive and accurate for complex biological interactions.

### Standardizing & annotating models

Reproducibility, a long-standing concern of the scientific community, can be significantly improved through the use of standards, annotations and repositories (32). The accurate and appropriate description of both the model itself and its components is also essential for model composition. As proposed by the curation and annotation of logical models (CALM) initiative (33), we use the Systems Biology Markup Language Qualitative Models (SBML-qual) format (34,35) to encode the selected models.

The community often refers to the HUGO Gene Nomenclature Committee (HGNC) guidelines for naming human genes (36). Additionally, the Vertebrate Gene Nomenclature Committee encourages the use of the human orthologs for other vertebrate species, such as for mouse and rat. To ensure accurate gene identifiers and facilitate the later composition process, we mapped each gene in the models to the HGNC gene ID and annotated them with the HGNC approved symbol. We implemented a semi-automated approach for this step, where standardized gene names can be queried through the HGNC REST API. After manual curation, the results can then be used to update the SBML-qual model. Following the International Protein Nomenclature Guidelines (37), we use the same abbreviations for proteins as their corresponding genes. Special attention should be given to fusion proteins and complexes, for which standard nomenclature may not exist. Metadata including original sources of the models should also be included in the annotation. We propose linking each component in the models to online resources, and annotating the supporting evidence of the edges, e.g., the experimental method used to determine the interaction, and the source of the data. This ensures that the models are transparent and compatible for integration.

### Reproducing selected models

As models become more complex, the reproducing task becomes more difficult (38,39). Before merging, each model is verified by replicating its published results under the same conditions. The results are then compared to the published performance to ensure that models are correctly implemented.

We used the CoLoMoTo Interactive Notebook in this study, which provides a unified environment for performing analyses and validating behavior of logical models (40). The advantage of using this tool is to ensure the reproducibility of results by using the same computational environment for different models.

### Composing models

Logical models represent GRNs as networks where nodes are genes or proteins, and edges represent regulatory interactions between them. The state of each node is determined by a logical rule *f*, which describes how the states of its regulators influence its activation. These logical functions are the core of the network and define its dynamic behavior. Using a combination of logical operations, we can then describe the updating schema for a gene or any biological molecule (41). We restrict the logical operators to logical product (AND, represented by the symbol ‘&’), logical sum (OR, symbol ‘|’), and logical negation (NOT, symbol ‘!’).

Suppose there are *n* logical models 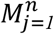 to be integrated, it is necessary to address both overlapping and non-overlapping components within them. For overlapping nodes that shared by the models, the logical rules *f*_*i*_ governing their behavior for each model *i* needs to be merged. We propose three deterministic methods for this merging process, each reflecting a different biological rationale:

#### 1. OR Combination: 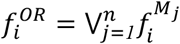

This method combines the logical rules from the individual models using the logical OR operator. If either of the rules from the models predicts the activation of a node, the combined rule will also predict activation. This approach ensures that the integrated model captures all possible activation scenarios, providing a more inclusive representation of the regulatory network.

#### 2. AND Combination: 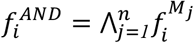

In contrast, the AND combination method uses the logical AND operator to merge the rules. Here, a node will only be activated in the integrated model if both original models predict its activation. This method is more stringent, ensuring that only consistent activation predictions are retained, which can reduce false positives and emphasize strong, corroborated regulatory interactions.

#### 3. Inhibitor Wins Combination: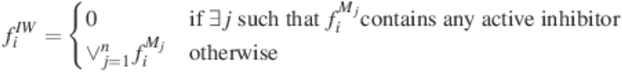

This method prioritizes inhibitory interactions. If any edge in the original models represents an inhibitory relationship, this inhibition will dominate in the merged model, leading to the node being turned off or having a negative impact on its status. This approach reflects the biological reality where inhibitory signals often have a strong regulatory effect, such as in the suppression of oncogenes or other critical pathways (7,42).

Given that gene regulation can be viewed as the interplay of transcription factors (TFs) and TF-binding sites of mRNA to govern expression levels of mRNA and their resulted proteins (43), it is intrinsically related to enhancer function and logic (44,45). Here, the three merging strategies offer complementary interpretations: the “OR” combination reflects the flexibility and robustness seen in enhancer integration, where multiple transcription factors (TFs) can independently activate a gene. The “Inhibitor Wins” approach represents the biological reality where repressive signals prevent inappropriate gene expression. This is supported by studies on transcriptional repression, where inhibitors can override activatory signals (46). The “AND” method enforces the strictest criteria, and is analogous to the cooperative binding of TFs for fine-tuned gene expression control. Researchers can select the most appropriate merging method based on the goals of the study, whether emphasizing inclusivity, stringency, or regulatory dominance. (See Supplementary data for an example.)

### Evaluating the merged model

Typically, evaluation of the logical models includes both dynamic and static analysis (47). Attractors or stable states can be associated with cellular phenotypes and represent the long-term behavior of the system. Simulation of the models can provide insights on possible trajectories, and can be used to predict response of the system to perturbations such as gene mutations. However, the specific methods should be determined by the user and the biological goal of the models. We provide some helper functions in the workflow for these tasks.

The aim of evaluation should be twofold: (1) to ensure that the integration maintains the functionality or performance of the original models, and (2) to demonstrate or explore new capabilities on new datasets. The first step is to verify if the merged model behaves similarly and retains the predictive accuracy of the individual models. Second, we test the merged model on new tasks to assess its robustness on different biological scenarios, such as predicting gene expression under new conditions or the impact of novel mutations. Finally, by comparing results from different integration strategies, we determine which approach best captures the biological processes and provides the most accurate predictions. We can then choose to employ the integration strategy that offers the highest accuracy and predictive power while aligning with the biological context of the study.

## Results: Use-case demonstration with AML models

We applied the workflow on two pairs of published logical models on AML to demonstrate the benefits of model integration for complex biological processes. A broad literature search was carried to identify relevant logical models related to gene regulation in AML. See Table S1 and S2 for the keywords and results. Because AML originates from hematopoietic stem cells (HSC) that have undergone malignant transformation (48), our search included models that describe HSC behavior. From the retrieved models, we selected two pairs of models based on their overlap and biological relevance. The first pair uses mouse data and investigates HSC development; the second pair uses human data and investigates AML disease progression. Details of the methods are provided in Supplementary data.

### Model pair 1: hematopoiesis

The first model pair consists of Boolean models constructed by Bonzanni et al. (49) and Krumsiek et al. (50) (Fig 2A). The former model captures key regulatory genes of early HSCs and simulates the differentiation of stem cells into mature blood cells, including erythroids, monocytes, and granulocytes. The latter one describes the transition of common myeloid progenitors into myeloid cells. The two models share 6 genes. Importantly, both models include key AML-related genes absent in the other model; the Bonzanni model includes *Runx1* while the Krumsiek model includes *Cebpa*, both of which are involved in the pathogenesis of AML (51,52). Merging these models results in a more detailed explanation of the HSCs differentiation process and thus possibly a more comprehensive representation for AML patients with different genetic alterations. (Note that this model pair was tested using mouse data, therefore, we use a different capitalization naming convention.)

**Fig 2:**
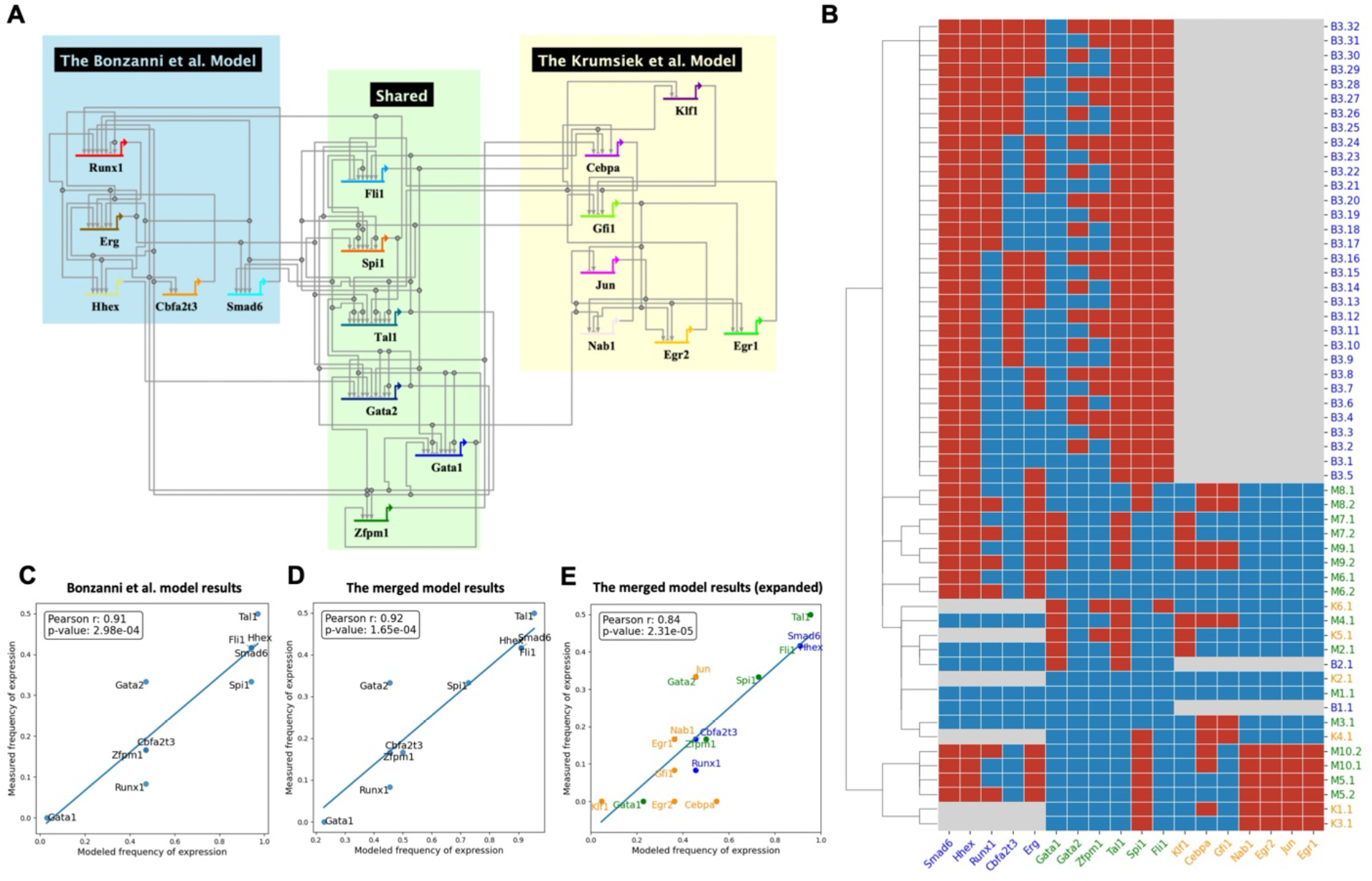
The merged model of Bonzanni et al. 2011 and Krumsiek et al. 2013 and its evaluation. (A) Visualization of the merged model using BioTapestry. Lines indicate regulation relationships that point from the regulator to its targets, with arrowheads as activating and bars as repressing. (B) Steady states pattern of the merged model pair and individual models. Each row is a steady state of the Bonzanni et al. model (start with ‘B’), Krumsiek et al. model (start with ‘K’), or the **merged model** (Start with ‘M’). The color in the heatmap indicates that a gene is ON (red), or OFF (blue), or that the gene is not included in the model (Grey). (C) Correlation of measured frequency of expression with the modeled frequency of activation for 10 genes using the Bonzanni et al. model. (D) Correlation of measured frequency of expression with the modeled frequency of activation for the 10 genes using the **merged model**. (E) Correlation of measured frequency of expression with the modeled frequency of activation for 18 genes using the **merged model**. 8 additional genes not covered in the Bonzanni et al. model are colored in orange. Results of the ‘OR’ model are shown, for other merged models results, see Fig S1 - S2.

We merged the models using our three different integration strategies: ‘AND’, ‘OR’ and ‘Inhibitor Wins’, and performed steady state analysis to identify possible attractors. Figure 2B shows the attractors for the simulation based on the ‘OR’ rule. To evaluate the consistency and robustness of the merged model, the attractor patterns of both the individual models and the merged model are clustered based on the hamming distance. A common goal of the two models is to describe cellular behavior of HSCs. It has been reported that each of the individual models can generate states that are representative of the different cell types that can originate from HSCs (49,50). Our results demonstrated the reproducibility of the original papers with the same stable states, with states B2.1 and K5.1 representative for the erythroid cells. One interesting finding is that the merged model combines them in a single state M2.1, where the expression of *Hhex, Runx1* and *Erg* are inactive and *Gata1, Tal1* and *Klf1* are active. This has been demonstrated by microarray experiments of erythroid cells (53). (See Fig S2 for the comparison.) Additionally, an interconnected cyclic attractor that consists of 32 states was identified in the original Bonzanni model, which represents a heterogeneous cell population of HSCs. The steady states derived from the merged model (M3.1 and M3.2) are clustered with them, featuring multiple genes active and *Gata1* consistently repressed. In contrast to the Bonzanni model, cyclic attractor M3.1 and M3.2 of the merged model do not show a variability in *Erg* and *Gata1*, suggesting their unique roles in HSC maintenance (54,55).

To evaluate the model quantitatively, we compared the frequency of expression of genes in stable states with gene expression data from single-cell microarray experiments of HSCs similar to the previous study (56). See Supplementary data for method details. Results of the merged model show a strong correlation (Fig 2D), comparable to the Bonzanni model alone (Fig 2C). We then asked whether the merged model is predictive for an extended set of genes from the Krumsiek model, and obtained a correlation of 0.84 (Fig 2E). Another observation is that the performance of merged models using different approaches varies, with the ‘OR’ model having the highest correlation (Fig S3). A possible reason is the heterogeneity of HSCs data, where multiple pathways may be activated simultaneously. This highlights the importance of choosing and evaluating approaches based on the context.

### Model pair 2: AML disease progression

Next, we evaluate a pair of models that focus on AML directly (Fig 3A). The Palma et al. model (7) integrates patient-specific genomic data into a Boolean network to predict clinical outcomes in AML patients, focusing on key regulatory pathways and frequent mutations in AML. The Ikonomi et al. model (57) uses a Boolean network to model HSC maintenance and the regulation of *TP53* pathway in response to niche interactions. These models overlap in key regulatory pathways involved in hematopoiesis and AML. By merging the two models, we can gain a deeper understanding of how genetic mutations in AML impact both the differentiation of blood cells and their interactions with signals from the stem cell environment, providing insights into the disease’s progression and potential therapeutic targets.

**Fig 3:**
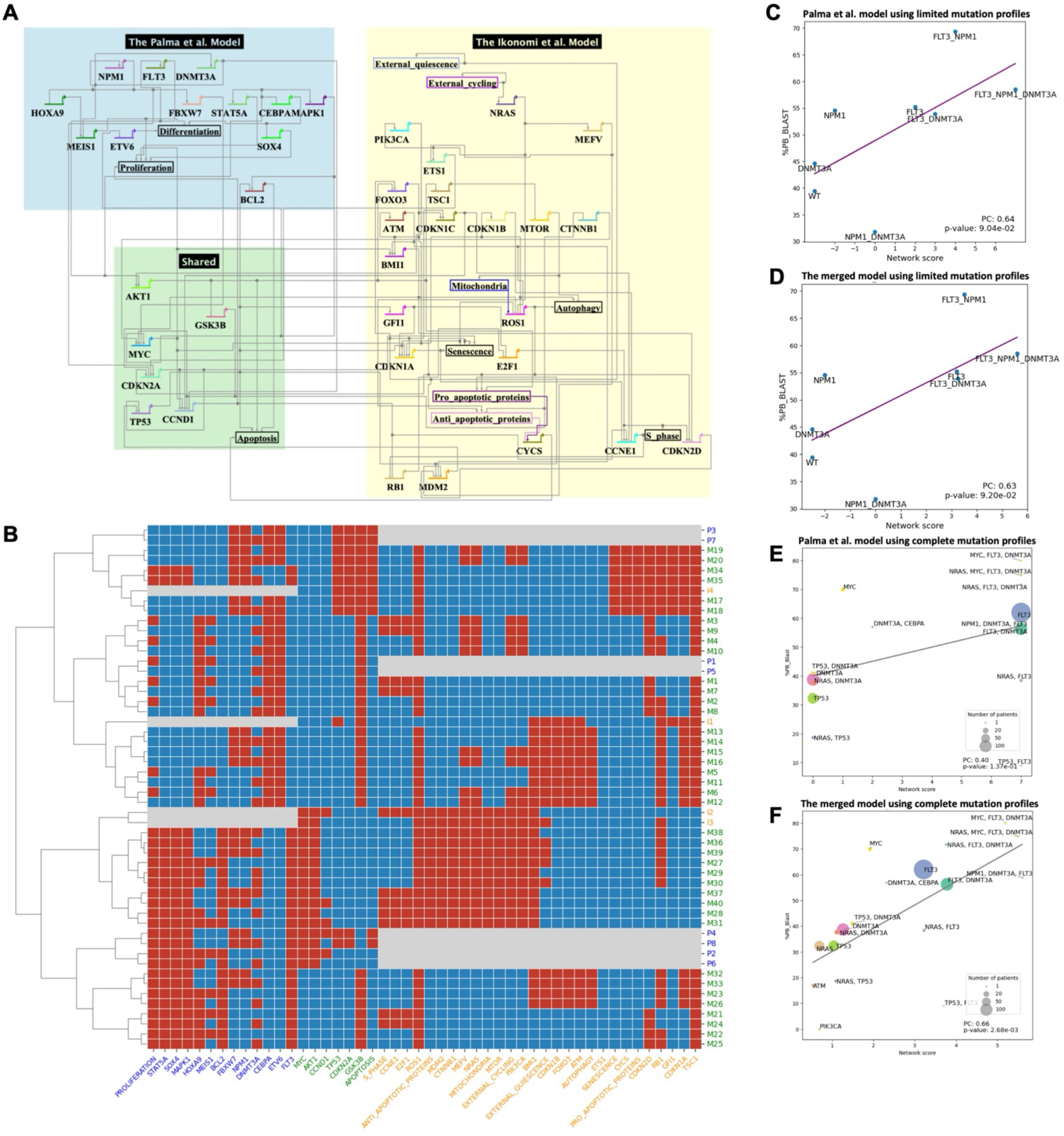
The merged model of Palma et al. 2021 and Ikonomi et al. 2020 and its evaluation. (A) Visualization of the merged model. (B) Steady states pattern of the **merged model** and individual models. (C) Correlation between the average blast percentage and network scores derived from the Palma et al. model for mutation status on FLT3, NPM1 and DNMT3A. (D) Correlation between blast percentage and network scores derived from the **merged model** for mutation status on FLT3, NPM1 and DNMT3A. (E) Correlation between blast percentage and network scores derived from the Palma et al. model for mutation status on all available genes in the Palma et al. model. Size of node indicates number of patients for each mutation profile. (F) Correlation between blast percentage and network scores derived from the **merged model** for mutation status on all available genes in the merged model. Only scatterplot of the ‘AND’ model is shown, for other merged models results, see Fig S4.

Similarly, the merged model replicates the steady states of the original models (Fig 3B). Each steady state from the individual models corresponds to several steady states in the merged model, which represents normal cellular function without perturbation. As an example, consider the attractors I1, from the Ikonomi model, and M13-M16, from the merged model. With external quiescence as input, these attractors represent the long-term state of normal HSCs where genes related to metabolism or cell growth are repressed (e.g., *MYC, SOX4, MAPK1, MTOR*), and cell cycle inhibitors are activated (e.g., *CDKN* genes, *RB1*). Furthermore, the merged model provides a more comprehensive view of gene regulation. Both original models only capture part of the regulations of the *TP53* tumor suppressor gene: the Palma model states that *ARF* (*CDKN2A*) activates *p53* (*TP53*); and the Ikonomi model describes the negative feedback of *MDM2*. Integrating the models provides a broader view of *p53* regulation, where *ARF* activates *p53* through the inhibition of *MDM2* (58).

An important modeling goal is to predict clinical outcomes for patients with specific mutations. For example, Palma et al. have connected their network to cancer hallmark phenotypes, and defined an integrated network score as a proxy for patients’ clinical outcomes (7). Similar to the original publication, we analyzed the correlation between the simulation outputs of the merged model and (1) the impact of mutations on overall survival, as measured by the Cox hazard ratio from the AMLSG dataset (23); and (2) the progression of disease, as measured by the percentage of blast cells from the TCGA dataset (17). Results suggest that the integration methods effectively capture the clinical implications of the genetic mutations (Table S3). Further, we applied the model to an independent big AML dataset, Beat AML (20). Both the Palma model (Fig 3C) and merged model (Fig 3D) show robustness of predicting disease progression with high correlation between the simulated scores from patients with gene mutations of *FLT3, NPM1* and *DNMT3A*, and the measured blast percentage.

One limitation is that patients with mutations in other genes, which may play an important role in AML prognosis, are not well modelled by a single model. Our goal with model merging is to capture a more comprehensive landscape, and improve the performance of prediction for a wider range of patients. Therefore, we expanded the patients’ mutation profiles to all available genes covered by the merged model. The performance of the Palma model declined when expanding the coverage, represented by the lower correlation between the model network score and clinical measurement of the blast percentage (Fig 3E), while the merged model improved the prediction power significantly (Fig 3F). This can be attributed to the richer information captured in the extended model. For example, clinical responses of the *TP53*-mutated patients can be modeled more accurately using the merged model, as shown by the light green circles. Since the Palma model, which only includes high-level interactions of the AML hallmark genes, the Ikonomi model compliments it by describing *TP53* regulation in more detail.

## Discussion

The study provides a semi-automatic approach to merge GRN models based on logic rules. One significant advantage for merging GRN models is to increase the coverage of patients with different mutations, thereby enhancing the model’s applicability. For example, *NRAS* mutated in 13.7% of the AML patients from the Beat AML dataset, but is not modelled by Palma et al. The potential impact of *NRAS* on disease progression can be described with improved accuracy using the merged Palma-Ikonomi model (Fig 3F). Similarly, merging the Bonzanni model with the Krumsiek model expands the coverage of 5.3% patients with *CEBPA* mutation. By including more genes in the models, they can better reflect the genetic diversity observed in AML patients.

There are different rules that can be defined for the merged models. Currently, users can select one of the three deterministic methods—AND, OR, or Inhibitor Wins—for the integration. The workflow allows to merge any number of models, although we have only tested it on pairs of models. While this provides a structured approach, it would be advantageous to establish a biologically-driven rationale for choosing among these methods. Furthermore, there may be instances where a single method applied across the entire network is insufficient. For example, a user might choose to apply the Inhibitor Wins method to nodes where inhibitory interactions dominate, such as genes with well-established repressive regulators, while using the AND method for nodes where activation requires strict synergy between inputs. As an alternative, one could also consider a probabilistic approach, where probabilities are assigned to each logical rule from the individual models (59,60). In biology, gene expression and TF binding have a stochastic nature, which is reflected in the random and context-dependent interactions among biological entities (61,62). Therefore, a probabilistic approach might better model the uncertainty and variability in regulatory interactions. In general, researchers will have to carefully consider how to merge rules from different models, depending on their research objectives and biological contexts. Model evaluation should be carried out to assess the performance of the models.

The application of this model merging workflow relies on existing models. The lack of comprehensive, high-quality models for certain genes and interactions in certain disease contexts became one obstacle. Although repositories for biosimulation models exist, they are primarily for internal validation and are not easily mapped to other external models (63). Moreover, extensive annotation is required to determine the biological relevance of the models, including their knowledge sources and the overall purpose of the model. One may consider merging models with similar contexts, and evaluate the merged model using new data. Automated model annotation is promising to advance the understanding and reuse of biosimulation models, and we plan to extend our previously built tool for quantitative SBML models (64) to qualitative models. The reproducibility of models is another critical aspect. In our study, we evaluated the reproducibility of each individual model before merging, and then compared the performance of the merged models with the original models across datasets. In the four models we considered, we were able to reproduce their performance (except for a minor discrepancy in the Palma model, which we corrected by contacting the authors). This highlights the importance of ensuring reproducibility to provide reliable predictions and support their applicability in broader research and clinical contexts. Finally, while the merging approaches are straightforward for Boolean models, they become more complex for multi-valued logical models, which would be a future step for this work, where the scales of the models are harmonized before composition.

In summary, we demonstrated the effectiveness of the workflow by merging two pairs of AML-focused models, capturing the complex interactions and regulatory mechanisms involved in gene expression and clinical outcomes. The merged models align well with biological phenomena and provide robust predictions, as evidenced by the strong correlations with experimental and clinical data. Our work also underscores the importance of standardization, reproducibility, and systematic documentation in logical models, and calls for comprehensive, high-quality data and extensive annotation. By addressing these challenges, the workflow has the potential to advance our understanding of complex biological systems and support the development of more effective and personalized therapies for diseases like AML.

Further biomedical applications may benefit from this work, such as constructing more comprehensive mechanistic models for diseases, which is one key aspect for biomedical digital twins (65). A digital twin is a set of virtual information constructs that mimics the structure, context, and behavior of the physical twin, and is dynamically updated with data from its physical twin (66). The model merging workflow allows researchers to systematically construct a virtual representation of a disease by integrating previously developed models. We argue that the merged models provide candidate models for biomedical digital twins, which in combination with pertinent patient data can be used to generate patient-specific models for predicting response to personalized treatments. In the case of AML, model personalization is possible by integrating mutation states of patients and drug response data available in datasets such as Beat AML. These personalized models could be used to test “*in silico*” for individual responses to drug interventions. Moving beyond AML, logical model merging could be applied in many other contexts wherever researchers have developed logical models of biological processes.

## Acknowledgements

This paper is dedicated to the memory of Dr. Ilya Shmulevich, the inventor of Probabilistic Boolean Networks in the early 2000s. Dr. Shmulevich is a former professor at Institute of Systems Biology, a leader of Cancer Genome Atlas Program (TCGA), and a pioneer in advancing cancer digital twin technology. As a great mentor and friend, Dr. Shmulevich has inspired all of us and many others in science and life.

## Author contributions

The work was initially proposed and conceptualized by G.Q. and J.G. L.L. developed the workflow, conducted data analyses, and wrote the initial draft of the manuscript. B.A. contributed to the conceptualization and methodological design of the study. G.Q. and J.G. provided supervision throughout the project. All authors reviewed, revised, and approved the final version of the manuscript.

## Data Availability

Models are collected from their publications. The individual model files and merged model files can be found at https://github.com/IlyaLab/LogicModelMerger/Models. Data used to test the models are all obtained from public repositories or publications, for details, please see Data section in the supplementary file.

## Reference

1. Karlebach G, Shamir R. Modelling and analysis of gene regulatory networks. Nat Rev Mol Cell Biol. 2008 Oct;9(10):770–80.

2. Le Novère N. Quantitative and logic modelling of molecular and gene networks. Nat Rev Genet. 2015 Mar;16(3):146–58.

3. Rothenberg EV. Causal Gene Regulatory Network Modeling and Genomics: Second-Generation Challenges. J Comput Biol. 2019 Jul 1;26(7):703–18.

4. Abou-Jaoudé W, Traynard P, Monteiro PT, Saez-Rodriguez J, Helikar T, Thieffry D, et al. Logical Modeling and Dynamical Analysis of Cellular Networks. Front Genet. 2016 May 31;7:94.

5. Sgariglia D, Carneiro FRG, Vidal de Carvalho LA, Pedreira CE, Carels N, da Silva FAB. Optimizing therapeutic targets for breast cancer using boolean network models. Comput Biol Chem. 2024 Apr;109:108022.

6. Plaugher D, Aguilar B, Murrugarra D. Uncovering potential interventions for pancreatic cancer patients via mathematical modeling. J Theor Biol. 2022 Sep 7;548:111197.

7. Palma A, Iannuccelli M, Rozzo I, Licata L, Perfetto L, Massacci G, et al. Integrating Patient-Specific Information into Logic Models of Complex Diseases: Application to Acute Myeloid Leukemia. J Pers Med. 2021 Feb 10;11(2):117.

8. Levine M, Davidson EH. Gene regulatory networks for development. Proc Natl Acad Sci. 2005 Apr 5;102(14):4936–42.

9. Davidson EH. Emerging properties of animal gene regulatory networks. Nature. 2010 Dec;468(7326):911–20.

10. Agmon E, Spangler RK, Skalnik CJ, Poole W, Peirce SM, Morrison JH, et al. Vivarium: an interface and engine for integrative multiscale modeling in computational biology. Bioinformatics. 2022 Mar 28;38(7):1972–9.

11. Liu ZP, Wu C, Miao H, Wu H. RegNetwork: an integrated database of transcriptional and posttranscriptional regulatory networks in human and mouse. Database J Biol Databases Curation. 2015 Sep 30;2015:bav095.

12. Pillich RT, Chen J, Churas C, Fong D, Gyori BM, Ideker T, et al. NDEx IQuery: a multi-method network gene set analysis leveraging the Network Data Exchange. Bioinformatics. 2023 Mar 1;39(3):btad118.

13. Karr JR, Takahashi K, Funahashi A. The principles of whole-cell modeling. Curr Opin Microbiol. 2015 Oct;27:18–24.

14. Karr JR, Sanghvi JC, Macklin DN, Gutschow MV, Jacobs JM, Bolival B, et al. A Whole-Cell Computational Model Predicts Phenotype from Genotype. Cell. 2012 Jul 20;150(2):389–401.

15. Macklin DN, Ahn-Horst TA, Choi H, Ruggero NA, Carrera J, Mason JC, et al. Simultaneous cross-evaluation of heterogeneous E. coli datasets via mechanistic simulation. Science. 2020 Jul 24;369(6502):eaav3751.

16. Georgouli K, Yeom JS, Blake RC, Navid A. Multi-scale models of whole cells: progress and challenges. Front Cell Dev Biol. 2023 Nov 7;11:1260507.

17. Cancer Genome Atlas Research Network, Ley TJ, Miller C, Ding L, Raphael BJ, Mungall AJ, et al. Genomic and epigenomic landscapes of adult de novo acute myeloid leukemia. N Engl J Med. 2013 May 30;368(22):2059–74.

18. Siveen KS, Uddin S, Mohammad RM. Targeting acute myeloid leukemia stem cell signaling by natural products. Mol Cancer. 2017 Jan 30;16(1):13.

19. Tyner JW, Tognon CE, Bottomly D, Wilmot B, Kurtz SE, Savage SL, et al. Functional genomic landscape of acute myeloid leukaemia. Nature. 2018 Oct;562(7728):526–31.

20. Bottomly D, Long N, Schultz AR, Kurtz SE, Tognon CE, Johnson K, et al. Integrative analysis of drug response and clinical outcome in acute myeloid leukemia. Cancer Cell. 2022 Aug 8;40(8):850-864.e9.

21. Malani D, Kumar A, Brück O, Kontro M, Yadav B, Hellesøy M, et al. Implementing a Functional Precision Medicine Tumor Board for Acute Myeloid Leukemia. Cancer Discov. 2022 Feb;12(2):388– 401.

22. Qin G, Dai J, Chien S, Martins TJ, Loera B, Nguyen QH, et al. Mutation Patterns Predict Drug Sensitivity in Acute Myeloid Leukemia. Clin Cancer Res Off J Am Assoc Cancer Res. 2024 Jun 14;30(12):2659–71.

23. Papaemmanuil E, Gerstung M, Bullinger L, Gaidzik VI, Paschka P, Roberts ND, et al. Genomic Classification and Prognosis in Acute Myeloid Leukemia. N Engl J Med. 2016 Jun 9;374(23):2209– 21.

24. Hérault L, Poplineau M, Duprez E, Remy É. A novel Boolean network inference strategy to model early hematopoiesis aging. Comput Struct Biotechnol J. 2023;21:21–33.

25. Silverbush D, Grosskurth S, Wang D, Powell F, Gottgens B, Dry J, et al. Cell-Specific Computational Modeling of the PIM Pathway in Acute Myeloid Leukemia. Cancer Res. 2017 Feb 15;77(4):827–38.

26. Helikar T, Kowal B, McClenathan S, Bruckner M, Rowley T, Madrahimov A, et al. The Cell Collective: toward an open and collaborative approach to systems biology. BMC Syst Biol. 2012 Aug 7;6:96.

27. Naldi A, Hernandez C, Abou-Jaoudé W, Monteiro PT, Chaouiya C, Thieffry D. Logical Modeling and Analysis of Cellular Regulatory Networks With GINsim 3.0. Front Physiol [Internet]. 2018 Jun 19 [cited 2024 May 21];9. Available from: https://www.frontiersin.org/journals/physiology/articles/10.3389/fphys.2018.00646/full

28. sybila/NewBioDiVinE [Internet]. sybila; 2014 [cited 2024 Jul 8]. Available from: https://github.com/sybila/NewBioDiVinE

29. Kanehisa M, Goto S. KEGG: kyoto encyclopedia of genes and genomes. Nucleic Acids Res. 2000 Jan 1;28(1):27–30.

30. Reactome Pathway Knowledgebase 2024 | Nucleic Acids Research | Oxford Academic [Internet]. [cited 2024 May 21]. Available from: https://academic.oup.com/nar/article/52/D1/D672/7369850?login=false&utm_source=advanceaccess&utm_campaign=nar&utm_medium=email

31. Lo Surdo P, Iannuccelli M, Contino S, Castagnoli L, Licata L, Cesareni G, et al. SIGNOR 3.0, the SIGnaling network open resource 3.0: 2022 update. Nucleic Acids Res. 2022 Oct 16;51(D1):D631–7.

32. Shin J, Porubsky V, Carothers J, Sauro HM. Standards, dissemination, and best practices in systems biology. Curr Opin Biotechnol. 2023 Jun 1;81:102922.

33. Niarakis A, Kuiper M, Ostaszewski M, Malik Sheriff RS, Casals-Casas C, Thieffry D, et al. Setting the basis of best practices and standards for curation and annotation of logical models in biology— highlights of the [BC]2 2019 CoLoMoTo/SysMod Workshop. Brief Bioinform. 2021 Mar 1;22(2):1848–59.

34. Chaouiya C, Bérenguier D, Keating SM, Naldi A, van Iersel MP, Rodriguez N, et al. SBML qualitative models: a model representation format and infrastructure to foster interactions between qualitative modelling formalisms and tools. BMC Syst Biol. 2013 Dec 10;7:135.

35. Chaouiya C, Keating SM, Berenguier D, Naldi A, Thieffry D, van Iersel MP, et al. The Systems Biology Markup Language (SBML) Level 3 Package: Qualitative Models, Version 1, Release 1. J Integr Bioinforma. 2015 Sep 4;12(2):270.

36. Seal RL, Braschi B, Gray K, Jones TEM, Tweedie S, Haim-Vilmovsky L, et al. Genenames.org: the HGNC resources in 2023. Nucleic Acids Res. 2023 Jan 6;51(D1):D1003–9.

37. International Protein Nomenclature Guidelines [Internet]. [cited 2024 Jun 4]. Available from: https://www.ncbi.nlm.nih.gov/genbank/internatprot_nomenguide/

38. Porubsky VL, Sauro HM. A Practical Guide to Reproducible Modeling for Biochemical Networks. In: Nguyen LK, editor. Computational Modeling of Signaling Networks [Internet]. New York, NY: Springer US; 2023 [cited 2024 Aug 7]. p. 107–38. Available from: 10.1007/978-1-0716-3008-2_5

39. Blinov ML, Gennari JH, Karr JR, Moraru II, Nickerson DP, Sauro HM. Practical resources for enhancing the reproducibility of mechanistic modeling in systems biology. Curr Opin Syst Biol. 2021 Sep 1;27:100350.

40. Naldi A, Hernandez C, Levy N, Stoll G, Monteiro PT, Chaouiya C, et al. The CoLoMoTo Interactive Notebook: Accessible and Reproducible Computational Analyses for Qualitative Biological Networks. Front Physiol. 2018 Jun 19;9:680.

41. Thomas R, D’Ari R. Biological Feedback. CRC Press; 1990. 328 p.

42. Deng X, Chen Y. Inference of Gene Regulations Between Multiple Activators/Inhibitors and Singular Genes. In: 2018 9th International Conference on Information Technology in Medicine and Education (ITME) [Internet]. 2018 [cited 2024 May 23]. p. 192–8. Available from: https://ieeexplore.ieee.org/document/8589283

43. Encyclopedia of Bioinformatics and Computational Biology: ABC of Bioinformatics. Elsevier; 2018. 3421 p.

44. Lambert SA, Jolma A, Campitelli LF, Das PK, Yin Y, Albu M, et al. The Human Transcription Factors. Cell. 2018 Feb 8;172(4):650–65.

45. Reiter F, Wienerroither S, Stark A. Combinatorial function of transcription factors and cofactors. Curr Opin Genet Dev. 2017 Apr 1;43:73–81.

46. Reynolds N, O’Shaughnessy A, Hendrich B. Transcriptional repressors: multifaceted regulators of gene expression. Development. 2013 Feb 1;140(3):505–12.

47. Samaga R, Klamt S. Modeling approaches for qualitative and semi-quantitative analysis of cellular signaling networks. Cell Commun Signal. 2013 Jun 26;11(1):43.

48. Long NA, Golla U, Sharma A, Claxton DF. Acute Myeloid Leukemia Stem Cells: Origin, Characteristics, and Clinical Implications. Stem Cell Rev Rep. 2022 Apr;18(4):1211–26.

49. Bonzanni N, Garg A, Feenstra KA, Schütte J, Kinston S, Miranda-Saavedra D, et al. Hard-wired heterogeneity in blood stem cells revealed using a dynamic regulatory network model. Bioinforma Oxf Engl. 2013 Jul 1;29(13):i80–88.

50. Krumsiek J, Marr C, Schroeder T, Theis FJ. Hierarchical differentiation of myeloid progenitors is encoded in the transcription factor network. PloS One. 2011;6(8):e22649.

51. Jalili M, Yaghmaie M, Ahmadvand M, Alimoghaddam K, Mousavi SA, Vaezi M, et al. Prognostic Value of RUNX1 Mutations in AML: A Meta-Analysis. Asian Pac J Cancer Prev APJCP. 2018 Feb 26;19(2):325–9.

52. Su L, Shi YY, Liu ZY, Gao SJ. Acute Myeloid Leukemia With CEBPA Mutations: Current Progress and Future Directions. Front Oncol. 2022 Feb 1;12:806137.

53. Chambers SM, Boles NC, Lin KYK, Tierney MP, Bowman TV, Bradfute SB, et al. Hematopoietic Fingerprints: An Expression Database of Stem Cells and Their Progeny. Cell Stem Cell. 2007 Nov 15;1(5):578–91.

54. Knudsen KJ, Rehn M, Hasemann MS, Rapin N, Bagger FO, Ohlsson E, et al. ERG promotes the maintenance of hematopoietic stem cells by restricting their differentiation. Genes Dev. 2015 Sep 15;29(18):1915–29.

55. Takai J, Moriguchi T, Suzuki M, Yu L, Ohneda K, Yamamoto M. The Gata1 5′ region harbors distinct cis-regulatory modules that direct gene activation in erythroid cells and gene inactivation in HSCs. Blood. 2013 Nov 14;122(20):3450–60.

56. Ramos CA, Bowman TA, Boles NC, Merchant AA, Zheng Y, Parra I, et al. Evidence for diversity in transcriptional profiles of single hematopoietic stem cells. PLoS Genet. 2006 Sep 29;2(9):e159.

57. Ikonomi N, Kühlwein SD, Schwab JD, Kestler HA. Awakening the HSC: Dynamic Modeling of HSC Maintenance Unravels Regulation of the TP53 Pathway and Quiescence. Front Physiol. 2020;11:848.

58. Sherr CJ, Weber JD. The ARF/p53 pathway. Curr Opin Genet Dev. 2000 Feb 1;10(1):94–9.

59. Shmulevich I, Dougherty ER, Kim S, Zhang W. Probabilistic Boolean networks: a rule-based uncertainty model for gene regulatory networks. Bioinformatics. 2002 Feb 1;18(2):261–74.

60. Trairatphisan P, Mizera A, Pang J, Tantar A, Schneider J, Sauter T. Recent development and biomedical applications of probabilistic Boolean networks. Cell Commun Signal. 2013;11(1):46.

61. McAdams HH, Arkin A. It’s a noisy business! Genetic regulation at the nanomolar scale. Trends Genet TIG. 1999 Feb;15(2):65–9.

62. Parab L, Pal S, Dhar R. Transcription factor binding process is the primary driver of noise in gene expression. PLOS Genet. 2022 Dec 12;18(12):e1010535.

63. Malik-Sheriff RS, Glont M, Nguyen TVN, Tiwari K, Roberts MG, Xavier A, et al. BioModels—15 years of sharing computational models in life science. Nucleic Acids Res. 2020 Jan 8;48(D1):D407– 15.

64. Shin W, Gennari JH, Hellerstein JL, Sauro HM. An automated model annotation system (AMAS) for SBML models. Bioinformatics. 2023 Nov 1;39(11):btad658.

65. Hernandez-Boussard T, Macklin P, Greenspan EJ, Gryshuk AL, Stahlberg E, Syeda-Mahmood T, et al. Digital twins for predictive oncology will be a paradigm shift for precision cancer care. Nat Med. 2021 Dec;27(12):2065–6.

66. Foundational Research Gaps and Future Directions for Digital Twins [Internet]. Washington, D.C.: National Academies Press; 2024 [cited 2024 Aug 26]. Available from: https://www.nap.edu/catalog/26894

