## Supplementary materials for "LM-Merger: A workflow for merging logical models with an application to gene regulation"

#### Merging Logical GRNs models

Mathematical models have been developed to capture the behavior of GRNs, among them, logical models are one of the simplest and most frequently used by biologists. This is partly due to limited information on kinetic parameters besides network structure, but primarily because logical models are intuitive and versatile (1,2). Despite their simplicity in formulation, logical models are able to generate and recapitulate complex behaviors often observed in cellular biology. Logical models use directed hypergraphs to describe not only pairwise relationships, as interaction graphs do, but also interactions between more than two components, which is common in biochemical reactions and signaling transductions (3). Since Kauffman's study in 1969 (4), logical models have been used to describe a wide range of activities in biological systems, from gene activities and protein expression to cellular behavior and system-level phenotypes, ultimately contributing to clinical and translational medicine (For reviews, see (2,5–7)).

Logical GRN models are represented as directed graph $G = (V, E)$, where $V$ is the set of nodes representing genes or proteins, and $E$ is the set of edges representing regulatory interactions. Each node $v_i \in V$ can take on a state $s_i$ from a finite set $S$, where $s_i \in S$. For Boolean models, $S = \{0,1\}$, where 0 represents an inactive (OFF) state and 1 represents an active (ON) state; and for multi-value logical models, $S = \{0,1, \dots, k\}$, where level of activation can be specified. Time is viewed as proceeding discretely in general logical models. States of the nodes can be updated synchronously, when all values are calculated after a transition, or asynchronously, with one at a time. At each step, the state of each node $v_i$ is determined by a logical rule $f_i$, which is a function of the states of its regulators. Formally, for each node $v_i$:

$$s_i(t+1) = f_i\big(s_1(t), s_2(t), \dots, s_n(t)\big)$$

where $s_i$ denotes the state of node $v_i$ at time $t$, $s_1$ through $s_n$ are the regulators of $s_i$, and $f_i$ is a logical function defined by logical operators.

The composition of multiple logical models involves identifying the overlapping and non-overlapping components of the models. Let $M_1, M_2, \dots, M_n$ be the models to be integrated. The sets of nodes and edges for each model are denoted as $(V_1, E_1)$ $(V_2, E_2), \dots, (V_n, E_n)$.

First, we identify the overlap $O$ and non-overlap $N_i$ components:

$$O = \bigcap_{i=1}^{n} V_i$$

$$N_i = V_i \setminus \bigcup_{j \neq i} V_j$$

For the overlapping components, the logical function $f_i$ for each node $v_i \in O$ needs to be updated. The merged logical function $f_i^{merged}$ is defined using logical combination methods. Here, we employ three methods, each with a unique rationale for gene or protein interactions:

1. **OR Combination**: $f_i^{OR} = \bigvee_{j=1}^{n} f_i^{M_j}$

   This method combines the logical rules from the individual models using the logical OR operator. If either of the rules from the models predicts the activation of a node, the combined rule will also predict activation. This approach ensures that the integrated model captures all possible activation scenarios, providing a more inclusive representation of the regulatory network.

2. **AND Combination**: $f_i^{AND} = \wedge_{j=1}^{n} f_i^{M_j}$

   In contrast, the AND combination method uses the logical AND operator to merge the rules. Here, a node will only be activated in the integrated model if both original models predict its activation. This method is more stringent, ensuring that only consistent activation predictions are retained, which can reduce false positives and emphasize strong, corroborated regulatory interactions.

3. **Inhibitor Wins Combination**: $f_i^{IW} = \begin{cases} 0 & \text{if } \exists j \text{ such that } f_i^{M_j} \text{contains any active inhibitor} \\ \bigwedge_{j=1}^{n} f_i^{M_j} & \text{otherwise} \end{cases}$

   This method prioritizes inhibitory interactions. If any edge in the original models represents an inhibitory relationship, this inhibition will dominate in the merged model, leading to the node being turned off or having a negative impact on its status. This approach reflects the biological reality where inhibitory signals often have a strong regulatory effect, such as in the suppression of oncogenes or other critical pathways (8,9).

By employing these combination methods, we can then select the one that best fits the goal of composition based on biological considerations and the specific requirements of the study (inclusivity, stringency, or regulatory dominance). For the non-overlapping components, the nodes and their associated logical functions are directly added to the merged model. The final merged model $M_{Merged}$ consists of:

$$V_{merged} = O \cup \bigcup_{i=1}^{n} N_i$$

$$E_{merged} = \bigcup_{i=1}^{n} E_i$$

The figure below shows an example of merging rules for a shared node, "*Apoptosis*," from two logical models on AML. In one model (9), apoptosis is regulated by activation of *TP53*, a tumor suppressor, and inhibition of *BCL2*, an anti-apoptotic protein. While another model (10) defines apoptosis as dependent on *CYCS* activation that facilitates cell death caspases, and *AKT1* inhibition that promotes tumor survival. Since these factors function through different mechanisms and pathways, the merging strategies offer different interpretations: the "OR" combination captures all possible conditions from both models; the "Inhibitor Wins" approach emphasizes inhibitory interactions by requiring at least one pro-apoptotic signal while excluding any anti-apoptotic influence; and the "AND" method enforces the strictest criteria, requiring all conditions from both models to be met. For the non-overlapping components, the nodes and associated logical functions can be directly added to the merged model. The final integrated model combines the rules and structures from all input models, providing a comprehensive and biologically relevant network. This approach enables researchers to select the most appropriate merging method based on the goals of the study, whether emphasizing inclusivity, stringency, or regulatory dominance.

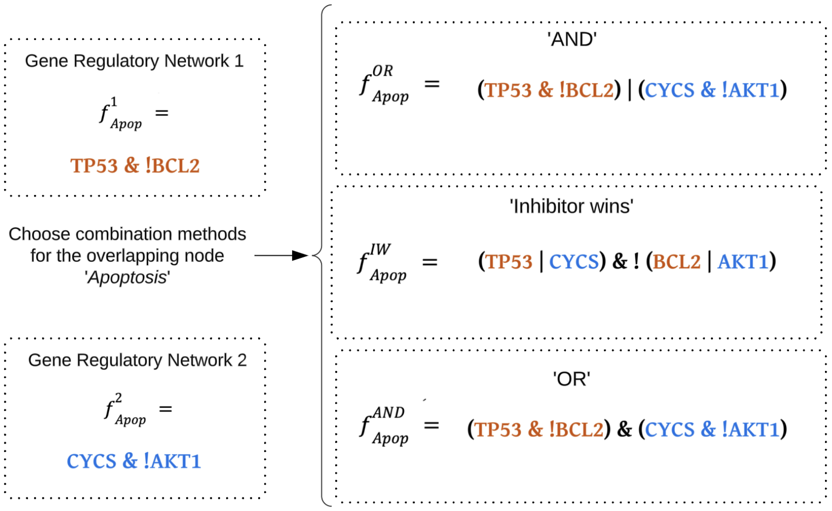


### Application to logical model composition for AML

We use AML-focused logical models to exemplify the model composition workflow, and evaluate performance of the merged models against original studies.

**Finding AML logical models**

We conducted a comprehensive literature search on PubMed to identify previously developed logical models related to gene regulation in AML. Specific search terms such as "AML gene regulatory network," "AML Boolean network model," and "AML qualitative network model" were used. The complete list of keywords used is provided in Table S1. Additional relevant papers were later identified through manual searches on Google Scholar and bioRxiv, addressing gaps not covered by the initial search terms. The initial search highlighted the need for manual review to identify truly relevant models, as studies sometimes use less specific terminology. For example, papers using terms like "signaling networks" or "signaling pathway" were not included, because adding those keywords result in more irrelevant papers on function of one specific gene or protein. Rigorous selection criteria were applied to ensure the relevance of the identified models. Models had to be AML-related and employ Boolean or qualitative logic. Preference was given to models validated with patient data or experimental evidence. Finally, the model pairs for merging should share an adequate level of overlapping genes.

Initially, 93 papers were retrieved; after review, we identified 19 of them that included a qualitative logical model and that were relevant to AML. Details of all 19 selected models, including model type, purpose of the model and validation information, are summarized in Table S2.

These 19 AML logical models share common purposes and validation approaches. Many rely on a combination of literature reviews, experimental data, and public knowledge resources or databases such as KEGG (11) and SIGNOR (12) for understanding molecular interactions and regulatory networks in AML. Most models use Boolean networks to describe interactions among genes and proteins, with some employing probabilistic Boolean models (13) or multi-valued models (14). Various computational tools, including GINsim (15), MaBoSS (16), BioModelAnalyzer (BMA) (17), Reactome FIViz (18), and caspo (19), are used for model construction and simulations. Validation methods primarily include in silico approaches, where the simulated behavior of the model is compared to previously generated experimental data or literature (used in 13 out of the 19 models). In vitro validation, involving laboratory testing of model predictions, is employed in 3 studies (20–22), while in vivo validation was not used in any of the models. Some models focus mainly on theoretical or computational aspects without extensive validation.

An important observation from our review is the lack of standardization and accessibility of these models, which leads to challenges in reproducibility. If model results are not reproducible, that greatly reduces their utility for further research. Our work underscores the need for improved standardization and accessibility, which is crucial for model reuse and model composition (23,24).

**Standardizing & annotating the AML logical models**

Next, we standardized these selected models into the SBML-qual format. Each model was annotated using the HGNC gene symbols to ensure accurate and consistent identifiers.

**Reproducing model results**

The reproducibility of these models is validated against their original publications using different computational tools including R, Python, and GINsim. Detailed methods and parameters used can be found in the original studies. Subsequent simulations and analyses were conducted in the CoLoMoTo Interactive Notebook using docker image colomoto/colomoto-docker:2024-03-01.

**Composing AML models**

Three approaches are tested for combining rules: the OR Combination, AND Combination, and Inhibitor Wins Combination as defined previously. The models are constructed manually at first, then compared with the automatically generated models to ensure consistency.

**Evaluating the merged models**

Steady state analysis using the asynchronous updating methods is performed for the Bonzanni - Krumsiek model pair; for the Palma - Ikonomi model pair, due to a larger network, synchronous updating is used. The expression patterns from the steady states of merged models are hierarchical clustered together with the steady states of individual models for comparison, and the nearest neighbor approach using Hamming distance was implemented for clustering. Further, the stable states from the Bonzanni - Krumsiek model pair are compared with gene expression data of blood cells generated by Chambers et al. (25). Data was obtained as GCRMA normalized microarray data. We calculated the average expression level of each cell type and converted them into dichotomous variables to compare with Boolean states.

To demonstrate benefits of the model composition, for the first model pair, we correlate the averaged gene expression activity from single-cell profiles of hematopoietic stem cells (HSCs) (26), as used in one of the original studies, with the average gene activity from modeled steady states in both individual and merged models. The modeled frequency of expression for each gene is calculated as the proportion of steady states in which the gene is 'on' (Boolean state of 1) relative to the total number of steady states in the model. For the second model pair, we identified patients' mutation profiles as the mutation status of three critical AML genes: *FLT3*, *NPM1* and *DNMT3A*. Phenotype scores were calculated for each profile as described by Palma et al.(9) and correlates with clinical outcomes: hazard ratio of death from the AMLSG dataset (27) and blast percentages from the TCGA-LAML dataset (28).

To cover the patients with other mutations, which could not be modeled using the previous approach, we tested the models' performance using all mutations available in the BeatAML dataset. For example, the Palma et al. model covers 18 genes, if a patient has any of them mutated, the combination of those genes would be used as his/her mutation profile. A similar approach was done for the merged model, except more genes are included. We identified the mutation profile of each patient using the mutation calls data and applied filters on the SIFT score (29) and PolyPhen score (30) to find only the deleterious variants. In addition, *FLT3-ITD* mutations derived from clinical genotyping results are added. The mutated tumor suppressor genes are set to a fixed value of 0 (loss of function), and the mutated oncogenes are set to a fixed value of 1 (gain of function) for subsequent simulation. For each profile, the phenotype scores are calculated as previously and compared with the average blast percentages of the patients.

#### Data

For individual models and the merged models used in this study, please see https://github.com/IlyaLab/LogicModelMerger/Models.

The following public data are used in the evaluation of the merged models:

1. Gene expression data of HSCs and their differentiated progeny using microarray analysis by Chambers et al. (25). Data was taken from Table S2. Complete GCRMA Normalized Hematopoietic Microarray Data Set. in the supporting information of the publication.

2. The averaged gene expression activity from single-cell profiles of HSCs provided by Ramos et al.(26). Data was taken from Table S6: MAS Calls in Each of the Samples Amplified with GSC RT-PCR Followed by Oligonucleotide Microarray Analysis in the supporting information of the publication.

3. Hazard ratio of death given by a clinical study of 1540 AML patients (AMLSG), where survival analysis was performed using Cox proportional-hazards methods(27). Mutation-specific hazards for *FLT3*, *NPM1*, *DNMT3A* and their combinations are given in Table 2 in the publication.

4. Blast percentages and genetic mutations of 200 AML patients from the TCGA-LAML study, where whole-genome sequencing (50 cases) or whole-exome sequencing (150 cases) are performed and clinical outcomes are collected(28). Mutation and clinical data were obtained from the NIH GDC website (https://gdc.cancer.gov/about-data/publications/laml_2012).

5. Blast percentages and genetic mutations of 805 AML patients from the Beat AML study, where a custom capture library (using whole-exome sequencing identified mutations) were used to generate the mutations of the patients. Mutation and clinical data were obtained from the BeatAML2 website (https://biodev.github.io/BeatAML2/).

### Supplementary tables and figures

Table S1: Keywords used in literature retrieval for AML-related logic models.

Keywords in the same category are combined using OR, and the two categories are combined using AND.

| Category | Keywords |
|---|---|
| Disease Related | Acute Myeloid Leukemia<br>AML<br>Myeloid<br>Hematopoiesis<br>Hematopoietic |
| Modeling Methods | Network Model<br>Boolean Network<br>Qualitative Network<br>Logic Model<br>Logical Model<br>Boolean Model<br>Qualitative Model<br>Network Models<br>Boolean Networks<br>Qualitative Networks<br>Logic Models<br>Logical Models<br>Network Modeling<br>Logic Modeling<br>Logical Modeling<br>Boolean Modeling<br>Qualitative Modeling |

Table S2: Summary of the identified AML-related logic models.

| DOI | Model Type | Topic | Purpose | Knowledge Source | Verification | Availability |
|---|---|---|---|---|---|---|
| 10.1073/pnas.1610622114 | Boolean network | myeloid and lymphoid development | Understand cellular behavior | ChIP-seq data of myeloid and lymphoid cells from GEO + Literature review | In silico | SBML and GINsim model available |
| 10.3390/jpm11030193 | Probabilistic Boolean Model | FLT3-mutant AML | Drug response | RNA-seq data of MV4-11 cells under 6 drug treatment conditions including quizartinib and dexamethasone + databases interactions (SIGNOR, TRRUST, RegNetwork) | NA | Txt files of Probabilistic and deterministic Boolean networks are available |
| 10.1016/j.csbj.2022.10.040 | Boolean network | HSC aging | Understand cellular behavior | Single cell RNA-seq analysis of mouse HSC and HSPC + Previous model (PMID: 21853041) | In silico | Full description in Fig. |
| 10.1158/0008-5472.CAN-16-1578 | Multi-valued logic model | AML | Drug response | Reverse-phase protein array (RPPA) of AML cell lines + literature review | in silico & in vitro | Full description in text. |
| 10.3390/jpm11020117 | Boolean network | AML | Clinical outcome prediction | Literature review | In silico | Full description in text. |
| 10.1186/s12859-018-2034-4 | Boolean network | AML | Clinical outcome prediction | Proteomics dataset of the AML DREAM 9 challenge (RPPA of patients) + KEGG PKN | In silico | Full description in Fig. |
| 10.1038/s41467-022-33189-w | Multi-valued logic model | AML | Clinical outcome prediction | Literature review | In silico | Full description in text and JSON file of the model. |

| | | | | | | |
|---|---|---|---|---|---|---|
| 10.1371/journal.pone.0022649 | Boolean network | Myeloid differentiation | Understand cellular behavior | Literature review | In silico | Full description in text. |
| 10.1038/srep08190 | Multi-valued logic model | CML | Drug response | Literature review | In silico | Only Figs. Rules are not clear. |
| 10.1186/1471-2105-15-S7-S7 | Boolean network | Myeloid differentiation | Understand cellular behavior | Microarray data from 3 datasets of myeloid blood cell lineage at ArrayExpress | NA | Could not find the model in the article (only a brief description) |
| 10.1016/j.isci.2022.104951 | Probabilistic Boolean network | Transdifferentiation from progenitor B cell to monocytic | Understand cellular behavior | Single-cell RNA-seq of bone marrow aspirates of healthy donors | in silico | Could not find the model in the article |
| 10.1016/j.csbj.2021.09.012 | Boolean network | HSC aging | Understand cellular behavior | Single-cell RNA sequencing data of isolated peripheral blood long-term HSCs from human | NA | Model in GitHub codes. |
| 10.1007/978-1-4939-9224-9_11 | Boolean network | HSC differentiation | Understand cellular behavior | Single-cell qRT-PCR data from hematopoietic stem and progenitor cells of mouse | NA | NA |
| 10.1073/pnas.1610609114 | Boolean network | HSC differentiation | Understand cellular behavior | Single-cell qRT-PCR data from hematopoietic stem and progenitor cells of mouse | NA | Full description in text. |
| 10.1093/bioinformatics/btx736 | Boolean network | Granulocyte-monocyte precursor-derived cells | Understand cellular behavior | Literature review | in silico | SBML files and codes are available at GitHub. The model is publicly available at Cell Collective. |
| 10.1007/978-1-0716-2277-3_12 | Boolean network | HSC homeostatic maintenance | Understand cellular behavior | Literature review | NA | NA |
| 10.3389/fphys.2020.00848 | Boolean network | HSC homeostatic maintenance | Understand cellular behavior | Literature review | in silico | Full description in text. |
| 10.1093/bioinformatics/btt243 | Boolean network | Early blood stem cell development | Understand cellular behavior | Literature review | in silico & in vitro | Full description in text. |
| 10.7554/eLife.90532.2 | Boolean network | AML | Drug response | SIGNOR + phosphorylation levels of sentinel proteins under 16 different perturbation conditions in TKIs sensitive and resistant cells | in silico & in vitro | Model file and codes are available at GitHub. |

Table S3: Summary of the correlation with clinical outcomes using the AMLSG and TCGA-LAML data.

| | | **Palma** | **'OR'** | **'Inhibitors win'** | **'AND'** |
|---|---|---|---|---|---|
| **Hazard ratio for death** | Correlation | 0.73 | 0.73 | 0.66 | 0.73 |
| | P-value | 0.041 | 0.041 | 0.077 | 0.040 |
| **Blast percentage** | Correlation | 0.71 | 0.68 | 0.69 | 0.67 |
| | P-value | 0.050 | 0.063 | 0.057 | 0.067 |

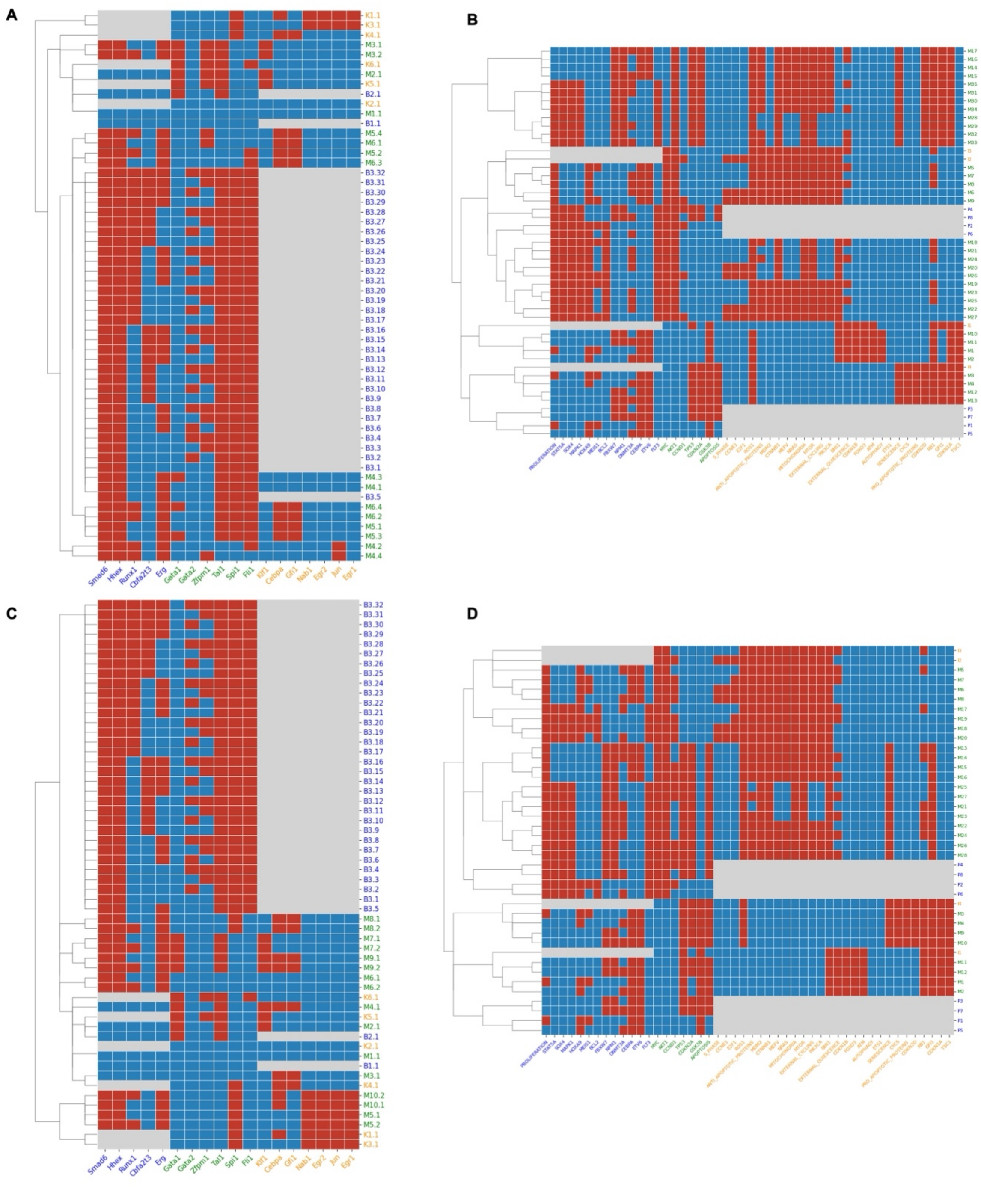


Fig S1: Steady states pattern of the merged models.

(A) Steady states pattern of the Bonzanni - Krumsiek merged model pair using the ‘Inhibitor wins’ approach. (B) Steady states pattern of the Palma - Ikonomi merged model pair using the ‘Inhibitor wins’ approach. (C) Steady states pattern of the Bonzanni - Krumsiek merged model pair using the ‘AND’ approach. (D) Steady states pattern of the Palma - Ikonomi merged model pair using the ‘OR’ approach. Steady states of both the individual models and merged models are calculated using the asynchronous approach and compared in the heatmap. The color in the heatmap indicates that a gene is ON (Red), OFF (Blue), or that the gene is not included in the model (Grey). For the axis labels, blue and orange indicate genes (x axis) or states (y axis) of the individual models, and green indicates genes shared by both models (x axis) or states of the merged model (y axis).

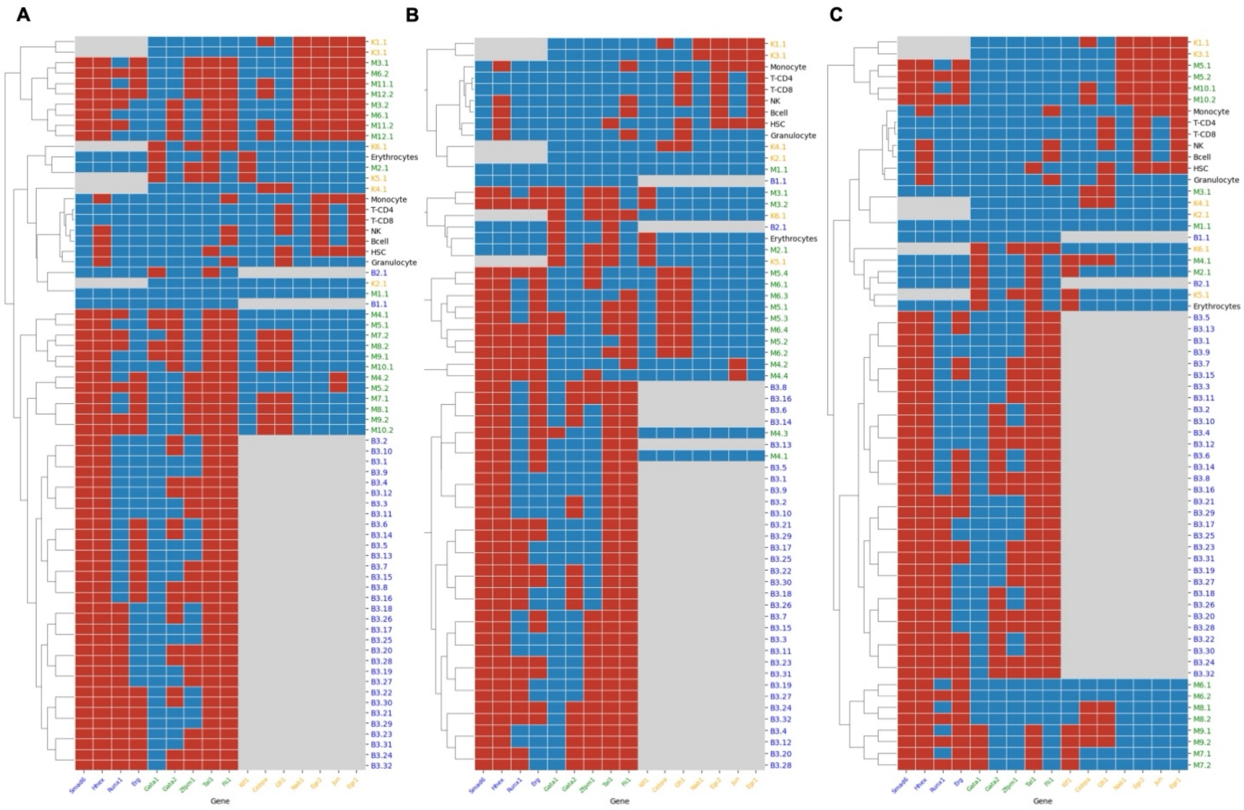


Fig S2: Compare steady states of the merged models with gene expression profiles of blood cells.

(A) Steady states of the Bonzanni - Krumsiek merged model pair using the ‘OR’ approach are clustered with gene expression of blood cells from Chambers et al. Color scales are similar as Figure S1, except that the black legends in the y-axis represent experimental data of each blood cell. (B) Steady states of the Bonzanni - Krumsiek merged model pair using the ‘Inhibitor wins’ approach (C) Steady states pattern of the Bonzanni - Krumsiek merged model pair using the ‘AND’ approach.

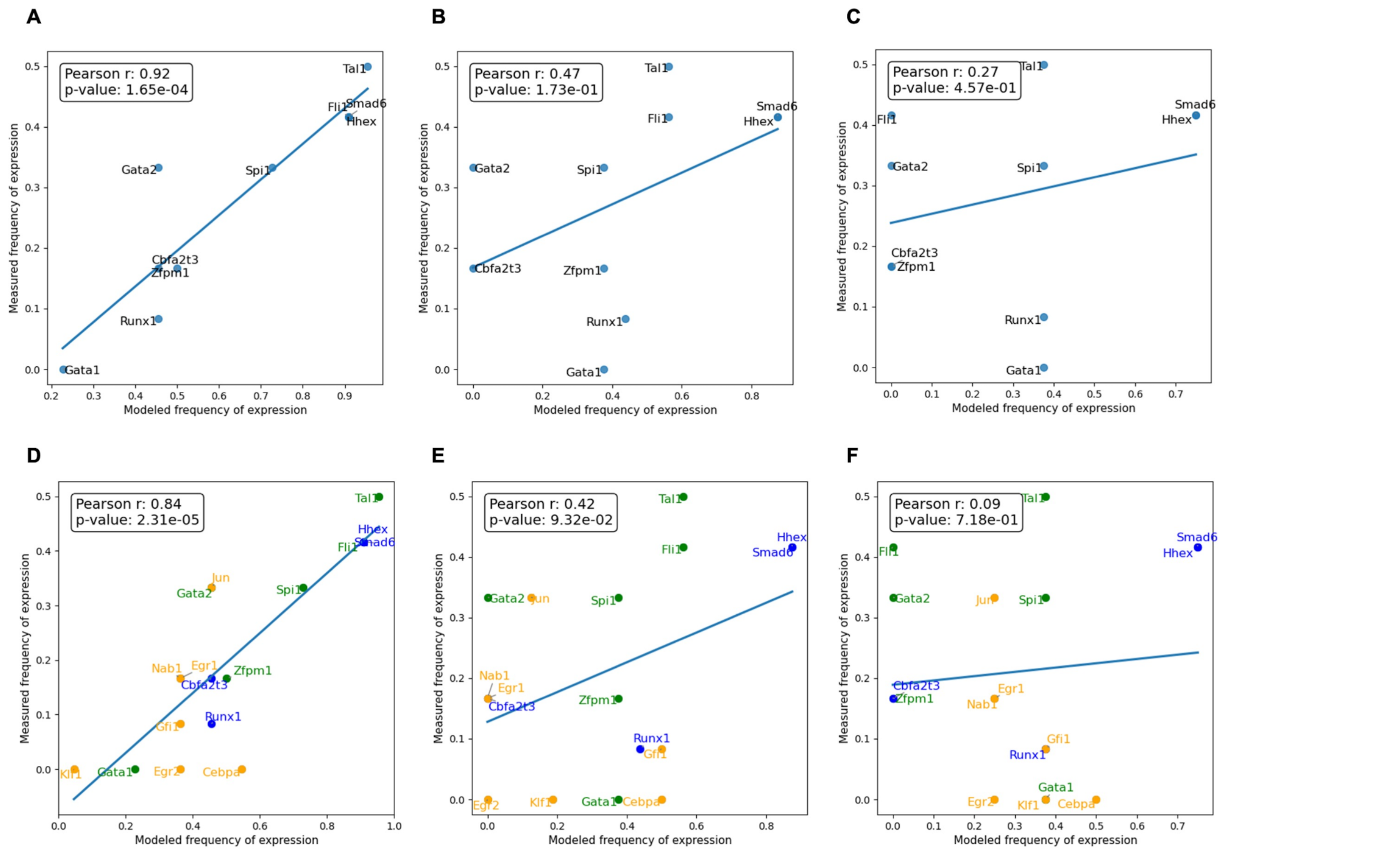


Fig S3: Expanding gene behavior modeling through merged models.

(A - C) Correlation of measured gene expression from the 12 single-cell profiles (Ramos et al., 2006) to the modeled frequency of activation for each of the 10 genes in the Bonzanni et al. model using the merged model of ‘OR’, ‘Inhibitor wins’ and ‘AND’ approach, respectively. (D - F) Correlation of the gene expression data with the modeled frequency of activation of 18 genes using the merged model of ‘OR’, ‘Inhibitor wins’ and ‘AND’ approach, respectively. 8 additional genes not covered in the Bonzanni et al. model are colored in orange, together with genes covered in the Bonzanni et al. model only (blue) and shared by both models (green).

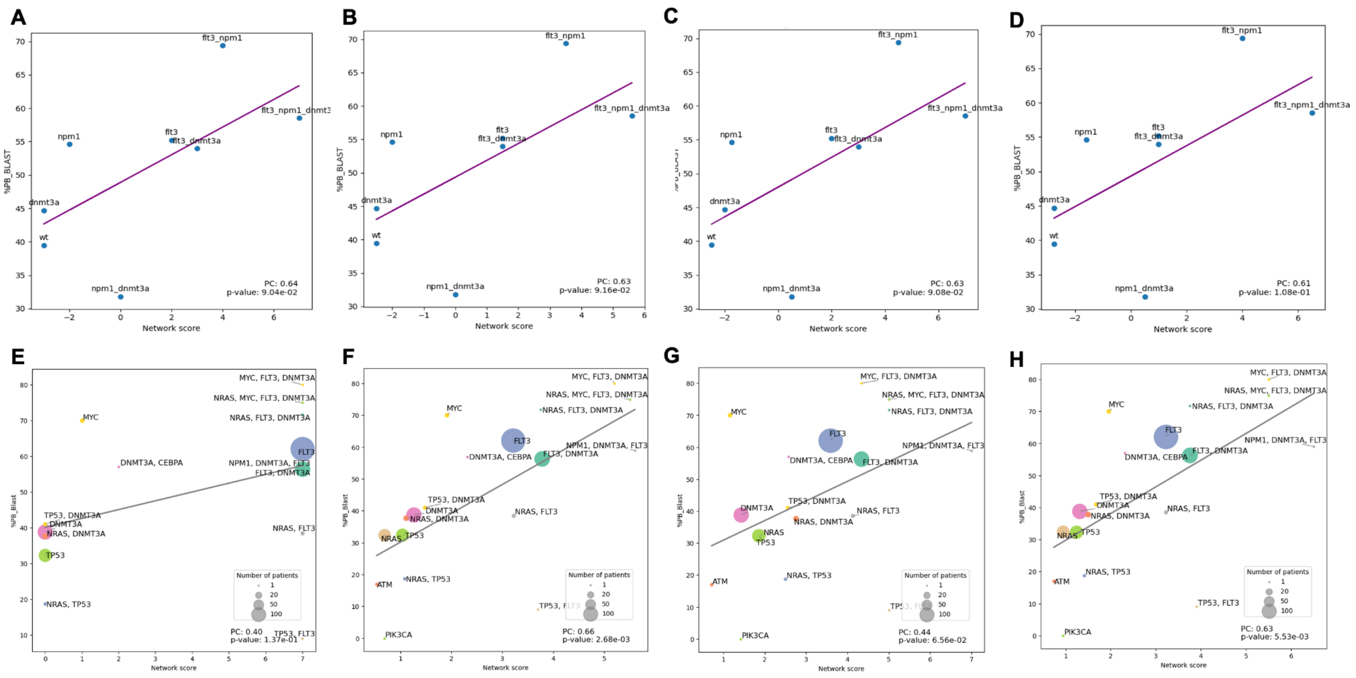


Fig S4: Expanding clinical outcome modeling through merged models.

Beat AML, a larger AML dataset, was used to evaluate the models' performance on a more diverse patient population. (A) Correlation between blast percentage of AML patients and network scores derived from the model for the Palma et al. model. (B-D) Correlation between blast percentage and network scores derived from the merged models using 'OR', 'Inhibitor wins' and 'AND' approach, respectively. (E) Instead of only three genes as tested previously, using all available genes in the Palma et al. model to predict blast percentage. Size of node indicates number of patients for each mutation profile. (F-H) Test on mutation status on all available genes using the merged models of 'OR', 'Inhibitor wins' and 'AND' approach, respectively.

#### Reference


1. Rothenberg EV. Causal Gene Regulatory Network Modeling and Genomics: Second-Generation Challenges. J Comput Biol. 2019 Jul 1;26(7):703–18.
2. Abou-Jaoudé W, Traynard P, Monteiro PT, Saez-Rodriguez J, Helikar T, Thieffry D, et al. Logical Modeling and Dynamical Analysis of Cellular Networks. Front Genet. 2016 May 31;7:94.
3. Samaga R, Klamt S. Modeling approaches for qualitative and semi-quantitative analysis of cellular signaling networks. Cell Commun Signal. 2013 Jun 26;11(1):43.
4. Kauffman SA. Metabolic stability and epigenesis in randomly constructed genetic nets. J Theor Biol. 1969 Mar;22(3):437–67.
5. Le Novère N. Quantitative and logic modelling of molecular and gene networks. Nat Rev Genet. 2015 Mar;16(3):146–58.
6. Traynard P, Tobalina L, Eduati F, Calzone L, Saez-Rodriguez J. Logic Modeling in Quantitative Systems Pharmacology: Logic Modeling in Quantitative Systems Pharmacology. CPT Pharmacomet Syst Pharmacol. 2017 Aug;6(8):499–511.
7. Hemedan AA, Niarakis A, Schneider R, Ostaszewski M. Boolean modelling as a logic-based dynamic approach in systems medicine. Comput Struct Biotechnol J. 2022 Jun 17;20:3161–72.
8. Deng X, Chen Y. Inference of Gene Regulations Between Multiple Activators/Inhibitors and Singular Genes. In: 2018 9th International Conference on Information Technology in Medicine and Education (ITME) [Internet]. 2018 [cited 2024 May 23]. p. 192–8. Available from: https://ieeexplore.ieee.org/document/8589283
9. Palma A, Iannuccelli M, Rozzo I, Licata L, Perfetto L, Massacci G, et al. Integrating Patient-Specific Information into Logic Models of Complex Diseases: Application to Acute Myeloid Leukemia. J Pers Med. 2021 Feb 10;11(2):117.
10. Ikonomi N, Kühlwein SD, Schwab JD, Kestler HA. Awakening the HSC: Dynamic Modeling of HSC Maintenance Unravels Regulation of the TP53 Pathway and Quiescence. Front Physiol. 2020;11:848.
11. Kanehisa M, Furumichi M, Sato Y, Kawashima M, Ishiguro-Watanabe M. KEGG for taxonomy-based analysis of pathways and genomes. Nucleic Acids Res. 2022 Oct 27;51(D1):D587–92.
12. Lo Surdo P, Iannuccelli M, Contino S, Castagnoli L, Licata L, Cesareni G, et al. SIGNOR 3.0, the SIGnaling network open resource 3.0: 2022 update. Nucleic Acids Res. 2022 Oct 16;51(D1):D631–7.
13. Shmulevich I, Dougherty ER, Kim S, Zhang W. Probabilistic Boolean networks: a rule-based uncertainty model for gene regulatory networks. Bioinformatics. 2002 Feb 1;18(2):261–74.
14. Thomas R, D'Ari R. Biological Feedback. CRC Press; 1990. 328 p.
15. Gonzalez AG, Naldi A, Sanchez L, Thieffry D, Chaouiya C. GINsim: a software suite for the qualitative modelling, simulation and analysis of regulatory networks. Biosystems. 2006;84(2):91–100.
16. Stoll G, Caron B, Viara E, Dugourd A, Zinovyev A, Naldi A, et al. MaBoSS 2.0: an environment for stochastic Boolean modeling. Bioinformatics. 2017;33(14):2226–8.
17. Benque D, Bourton S, Cockerton C, Cook B, Fisher J, Ishtiaq S, et al. Bma: Visual Tool for Modeling and Analyzing Biological Networks. In: Madhusudan P, Seshia SA, editors. Computer Aided Verification [Internet]. Berlin, Heidelberg: Springer Berlin Heidelberg; 2012 [cited 2024 Sep 13]. p. 686–92. (Hutchison D, Kanade T, Kittler J, Kleinberg JM, Mattern F, Mitchell JC, et al., editors. Lecture Notes in Computer Science; vol. 7358). Available from: http://link.springer.com/10.1007/978-3-642-31424-7_50
18. Wu G, Dawson E, Duong A, Haw R, Stein L. ReactomeFIViz: a Cytoscape app for pathway and network-based data analysis. F1000Research. 2014 Sep 12;3:146.
19. Videla S, Saez-Rodriguez J, Guziolowski C, Siegel A. caspo: a toolbox for automated reasoning on the response of logical signaling networks families. Bioinformatics. 2017;33(6):947–50.
20. Silverbush D, Grosskurth S, Wang D, Powell F, Gottgens B, Dry J, et al. Cell-Specific Computational Modeling of the PIM Pathway in Acute Myeloid Leukemia. Cancer Res. 2017 Feb

15;77(4):827–38.
21. Bonzanni N, Garg A, Feenstra KA, Schütte J, Kinston S, Miranda-Saavedra D, et al. Hard-wired heterogeneity in blood stem cells revealed using a dynamic regulatory network model. Bioinforma Oxf Engl. 2013 Jul 1;29(13):i80-88.
22. Latini S, Venafra V, Massacci G, Bica V, Graziosi S, Pugliese GM, et al. Unveiling the signaling network of FLT3-ITD AML improves drug sensitivity prediction. eLife [Internet]. 2023 Oct 23 [cited 2023 Dec 17];12. Available from: https://elifesciences.org/reviewed-preprints/90532
23. Porubsky VL, Sauro HM. A Practical Guide to Reproducible Modeling for Biochemical Networks. In: Nguyen LK, editor. Computational Modeling of Signaling Networks [Internet]. New York, NY: Springer US; 2023 [cited 2024 Aug 7]. p. 107–38. Available from: https://doi.org/10.1007/978-1-0716-3008-2_5
24. Blinov ML, Gennari JH, Karr JR, Moraru II, Nickerson DP, Sauro HM. Practical resources for enhancing the reproducibility of mechanistic modeling in systems biology. Curr Opin Syst Biol. 2021 Sep 1;27:100350.
25. Chambers SM, Boles NC, Lin KYK, Tierney MP, Bowman TV, Bradfute SB, et al. Hematopoietic Fingerprints: An Expression Database of Stem Cells and Their Progeny. Cell Stem Cell. 2007 Nov 15;1(5):578–91.
26. Ramos CA, Bowman TA, Boles NC, Merchant AA, Zheng Y, Parra I, et al. Evidence for diversity in transcriptional profiles of single hematopoietic stem cells. PLoS Genet. 2006 Sep 29;2(9):e159.
27. Papaemmanuil E, Gerstung M, Bullinger L, Gaidzik VI, Paschka P, Roberts ND, et al. Genomic Classification and Prognosis in Acute Myeloid Leukemia. N Engl J Med. 2016 Jun 9;374(23):2209–21.
28. Cancer Genome Atlas Research Network, Ley TJ, Miller C, Ding L, Raphael BJ, Mungall AJ, et al. Genomic and epigenomic landscapes of adult de novo acute myeloid leukemia. N Engl J Med. 2013 May 30;368(22):2059–74.
29. Ng PC, Henikoff S. SIFT: predicting amino acid changes that affect protein function. Nucleic Acids Res. 2003 Jul 1;31(13):3812–4.
30. Adzhubei I, Jordan DM, Sunyaev SR. Predicting Functional Effect of Human Missense Mutations Using PolyPhen-2. Curr Protoc Hum Genet Editor Board Jonathan Haines Al. 2013 Jan;0 7:Unit7.20.
